# Neuropilin-1 regulates the secondary CD8 T cell response to virus infection

**DOI:** 10.1101/605816

**Authors:** Ji Young Hwang, Yanbo Sun, Christopher R. Carroll, Edward J Usherwood

## Abstract

Neuropilin-1 plays important roles in axonal guidance in neurons, and in the growth of new blood vessels. There is also a growing appreciation for roles played by neuropilin-1 in the immune response. This molecule is important for the function of regulatory T cells, however roles in other T cell populations have not been identified. Here we show that neuropilin-1 is expressed during the peak of the antiviral CD8 T cell response during murine gammaherpesvirus infection. Using a conditional knockout model, we deleted Nrp1 either before infection, or after CD8 T cell memory had been established. We found deletion of Nrp1 skewed the acute CD8 T cell response toward a memory precursor-like phenotype, however the ensuing resting memory response was similar regardless of Nrp1 expression. Interestingly Nrp1 deletion had differing effects on the recall response depending on the timing of deletion. When deleted before infection, Nrp1 deficiency inhibited the secondary response. Deletion just prior to re-exposure to virus lead to an enhanced secondary response. Interestingly these effects were observed only in mice infected with a persistent strain of murine gammaherpesvirus, and not a non-persistent mutant strain. These data highlight a multifaceted role for neuropilin-1 in memory CD8 T cell differentiation, dependent upon the stage of the T cell response and characteristics of the infectious agent. Several therapeutic anti-cancer therapies focus on inhibition of Nrp1 to restrict tumor growth, so knowledge of how Nrp1 blockade may affect the CD8 T cell response will provide a better understanding of treatment consequences.

**Importance:** CD8 T cell responses are critical to control both virus infections and tumors. The ability of these cells to persist for long periods of time can result in lifelong immunity, as relatively small populations of cell can expand rapidly to counter re-exposure to the same insult. Understanding the molecules necessary for this rapid secondary expansion is critical if we are to develop therapies that can provide lifelong protection. This report shows an important and complex role for the molecule neuropilin-1 in the secondary response. Several cancer therapies targeting neuropilin-1 are in development, and this work will lead to better understanding of the effect these therapies could have upon the protective CD8 T cell response.

## INTRODUCTION

Neuropilin-1 (Nrp1) is a type I transmembrane protein with multiple domains that functions as a co-receptor for several ligands, such as semaphorins (SEMA), vascular endothelial growth factor (VEGF), and transforming growth factor beta (TGFβ). The molecule itself lacks kinase activity, but it associates with other receptors such as integrins, plexins and VEGF receptor, that mediate transmembrane signaling^1^. It was first studied in the nervous system, where neuropilin-1 is known to participate in neuronal development and provide cues for axonal guidance^2^. Later the interaction between VEGF and Nrp1 was found to play an important role in angiogenesis^3, 4^. The involvement of Nrp1 in the growth of new blood vessels in tumor vasculature promotes tumor progression, and its blockade can restrict tumor growth^5, 6^. Nrp1 can also be expressed by tumor cells themselves, and a peptide which inhibits VEGF-Nrp1 interactions has been shown to induce a apoptosis of Nrp1-expressing breast tumor cells^7^.

Tumors often elaborate an immunosuppressive microenvironment, and neuropilin-1 has been shown to play important roles in suppression mediated by regulatory T cells (T_reg_). A recent study showed Nrp1 on T_reg_ was required for the suppression of the anti-tumor T cell response, and to cure inflammatory colitis^8^. Engagement of Nrp1 promoted T_reg_ quiescence and limited differentiation, resulting in enhanced T_reg_ stability in the tumor^8^. Expression of Nrp1 differentiates natural from inducible regulatory T cells in some physiological settings^9, 10^, and also identifies CD4^+^C25^−^ T cells with inhibitory function and the ability to recruit conventional T_reg_^11^. Interestingly a recent report showed Nrp1 expression on group 3 innate lymphoid cells (ILC3s) in the lung with lymphoid tissue inducer activity, and suggested functions in the early development of tertiary lymphoid aggregates in the lung and/or pulmonary angiogenesis^12^. Collectively these studies imply Nrp1 not only has a major impact in modulating responses to tumors, but also plays a role in immune regulation and tissue remodeling other physiological settings.

Nrp1 clearly plays important roles in immune regulatory cell populations, however its role on conventional T cell populations has not been determined. CD8 T cells are important in controlling virus infections and restricting growth of tumors, providing lifelong immunity by developing into memory cells that can respond rapidly to re-infection. Memory cells develop from memory precursors present early in the T cell response^13^, and signals from cytokines, costimulatory molecules and CD4 T cells are necessary for them to develop optimal recall responses^14^. In this study, we investigated the role of Nrp1 on the CD8 T cell response to murine gammaherpesvirus (MHV-68) infection using a conditional knockout model. By means of this strain, we could restrict the deletion only to CD8 T cells, and regulate the timing of the deletion by tamoxifen administration.

We show that Nrp1 is highly upregulated on CD8 T cells during the acute phase of viral infection, and deletion of Nrp1 during this window skewed the T cells more toward memory precursors than terminally differentiated effector cells. Interestingly, ‘early’ Nrp1 deletion resulted in weaker CD8 T cell expansion following virus rechallenge, suggesting Nrp1 signaling during priming promotes optimal ‘programming’ of memory CD8 T cells. Interestingly when deletion of Nrp1 occurred just before the recall response, the magnitude of the response was higher, indicating Nrp1 signals restrain the recall response. Interestingly these effects were only observed with a persistent strain of MHV-68, but Nrp1 did not appear to affect recall responses in infection with a non-persistent strain of the virus. What emerges is a complex role for Nrp-1 in the CD8 T cell response, which is dependent both upon the timing of Nrp-1 expression during the primary vs secondary response, and the nature of the infection.

## MATERIALS AND METHODS

### Mice and MHV-68 infection

C57BL/6NCrl (B6) mice were originally obtained from Charles River Laboratory. Nrp1 E8i-CreERT2 R26-YFP mice were kindly provided by Dr. Dario A. Vignali, from The University of Pittsburgh. Primers used to genotype the strain were as follows:

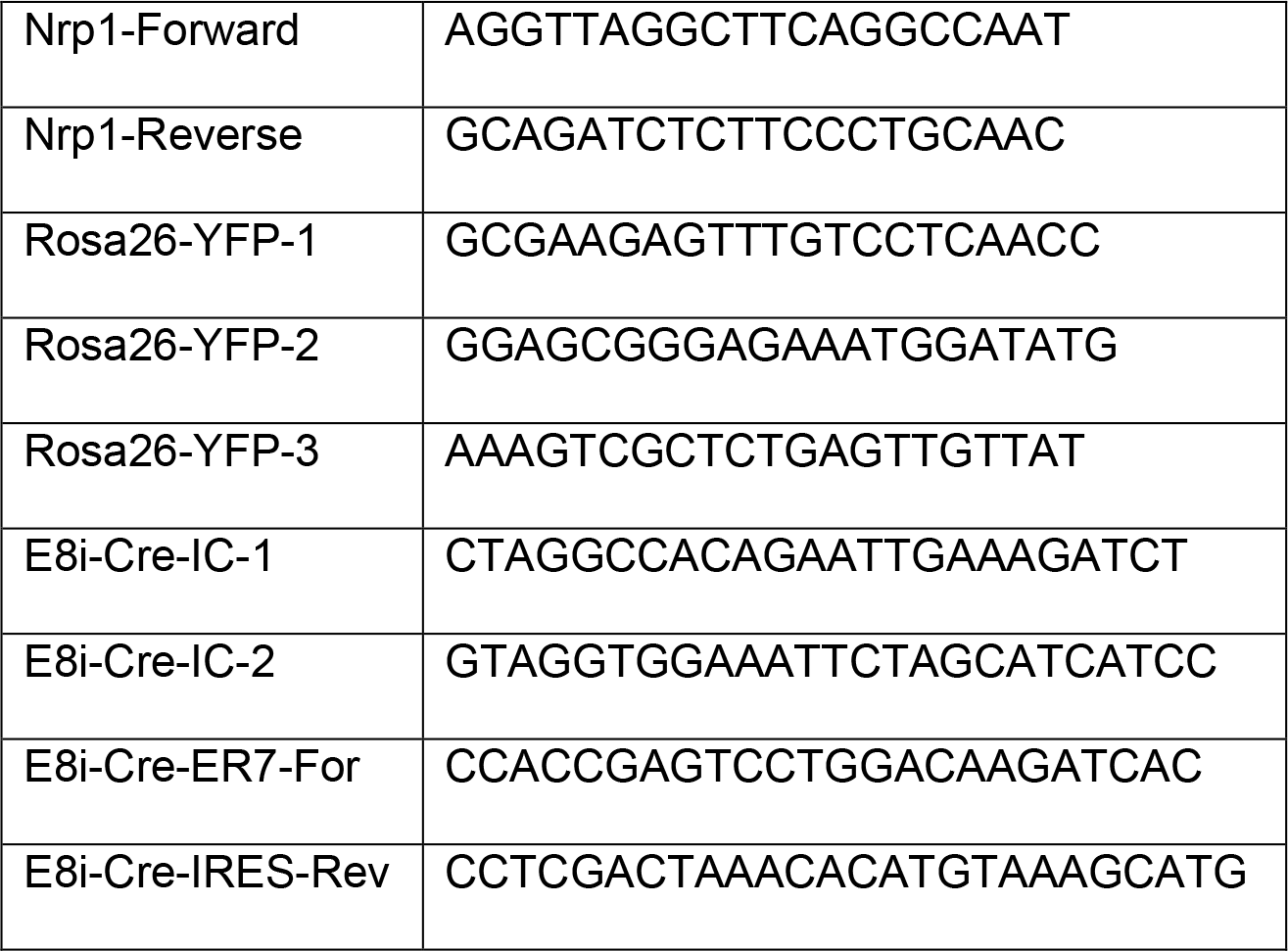

Mice were maintained under specific pathogen–free conditions in the Dartmouth Center for Comparative Medicine and Research. The Animal Care and Use Committee of Dartmouth College approved all animal experiments. MHV-68 containing a frameshift mutation in ORF73 (FS73) and the revertant virus (FS73R) were originally obtained from Stacey Efstathiou at The University of Cambridge, UK. 7-week-old mice were primarily infected with 4×10^3^ PFU by the intranasal route in 30μl Hank’s balanced salt solution (HBSS), and rechallenged with 1×10^6^ PFU WT MHV-68 by the intraperitoneal route.

### Tamoxifen treatment, T cell purification, and adoptive transfer

Tamoxifen (VWR) was suspended in 5% (v/v) EtOH-corn oil (Ward’s science), warmed at 37°C for at least 30 minutes before treatment, and 1 mg/100μl/mouse was given intraperitoneally for 5 consecutive days; starting at day −6 relative to infection (d-6) for early Nrp1 deletion and d28 for late Nrp1 deletion. CD8 T cells were purified from the spleens of early Nrp1 deleted (d28) or late Nrp1 deleted (d34) Nrp1 E8i-CreERT2 R26-mice using EasySep™ Mouse CD8 T Cell Isolation Kits (Stemcell Technologies) according to the manufacturer’s instructions. Single T cell preparations were >95% pure as determined by flow cytometry. CD8 T cell populations containing 2×10^4^ ORF61 tetramer^+^ memory CD8 T cells were injected by the retro-orbital route into B6-Ly5.1 recipients. The recipients were rechallenged one day after the adoptive T cell transfer, and splenocytes were collected on day 6 post-infection.

### Cell preparation, flow cytometry, and proliferation assay

Single-cell suspensions from spleen were prepared by passing them through cell strainers, and resuspended in Gey’s solution (150 mM NH_4_Cl, 10 mM KHCO_3_, and 0.05% phenol red) for 5 min to lyze red cells. Cell suspensions were then filtered through a 70 µm nylon cell strainer (BD Biosciences), washed, and resuspended in PBS with 2% bovine growth serum (BGS) and APC-conjugated tetramer specific for the MHV-68 dominant epitope (ORF61, NIH tetramer core facility) at room temperature for 1 hour, followed by 10 µg/ml Fc Block (2.4G2; Dartab) on ice for 10 min before staining with the following fluorochrome-conjugated antibodies (Abs): anti-cluster of differentiation (CD) 8-BV510 (CD8; 53-6.7), anti-CD45.2-BV421 (104), anti-CD45.2-BV650 (104), anti-CD304-BV421 (Neuropilin-1; 3E12), and anti-KLRG1-PE-Cy7 (2F1/KLRG1; all from BioLegend), and anti-CD4-APC (GK.15) and anti-CD127-APC 780 (A7R34; all from eBioscience). LIVE/DEAD™ Fixable Near-IR Dead Cell Stain Kit (Thermo Fisher) was used of cell viability, and Click-iT™ Plus EdU Pacific Blue™ Flow Cytometry Assay kit (Thermo Fisher) was used to assess cell proliferation. Cells were analyzed with MACSQuant (Miltenyi) FACS Aria (Becton Dickinson) or CytoFLEX S (Beckman Coulter) flow cytometers at the Dartlab flow cytometry core facility.

### Statistical Analysis

Two way ANOVA-Sidak’s or Dunnett’s multiple comparison test was used (GraphPad Prism Version 7.0). P values of less than 0.05 were considered statistically significant.

## RESULTS

### Nrp1 is upregulated with both persistent and non-persistent MHV-68 infection

Our previous research has shown the CD8 T cell response differs between infection with a mutant MHV-68 with a deletion in ORF73 (FS73)^15, 16^, which is essential for latent infection, when compared with a revertant virus that retains the ability to persist in the host. In order to understand the role of Nrp1 on CD8 T cells upon MHV-68 infection, we initially measured the kinetics of Nrp1 expression on CD8 T cells after either persistent (FS73R) or non-persistent (FS73) MHV-68 infection. Mice were infected with the relevant virus, then at various times post infection spleens cells were stained with MHC/peptide tetramers and anti-CD8 antibody to measure the frequency of CD8 T cells recognizing the dominant epitope^17^ (Fig. 1A and B). Consistent with our previous studies^16^, the magnitude of the CD8 T cell response was greater in the FS73R infected mice during the first four weeks of infection, however memory populations were of similar size in both strains (Figs 1A and B). Nrp1 expression was low in both cases during the early stages of infection (d7), but were significantly upregulated on d14, when CD8 T cell responses peak in MHV-68 infection^16,17^ (Fig. 1C). Nrp-1 expression slowly declined after 14 days, and had reduced to baseline expression levels by 60 days post-infection. While Nrp-1 was induced with these kinetics in both FS73 and FS73R infection, the induction was significantly greater after FS73 infection from days 14-21 post infection, but not significantly different thereafter (Fig. 1C and D). This lead to the T cell response to FS73 being dominated by Nrp1^hi^ cells during the acute infection (Fig. 1E), whereas there were more similar proportions of Nrp1^hi^ and Nrp1^lo^ cells at most times during the response to FS73R (Fig. 1F). In both cases the majority of memory CD8 T cells at d100 were Nrp1^hi^ (Figs 1E and 1F). These data indicate the absence of persistent infection leads to a greater induction of Nrp1 in the responding CD8 T cell population.

**Figure 1.**
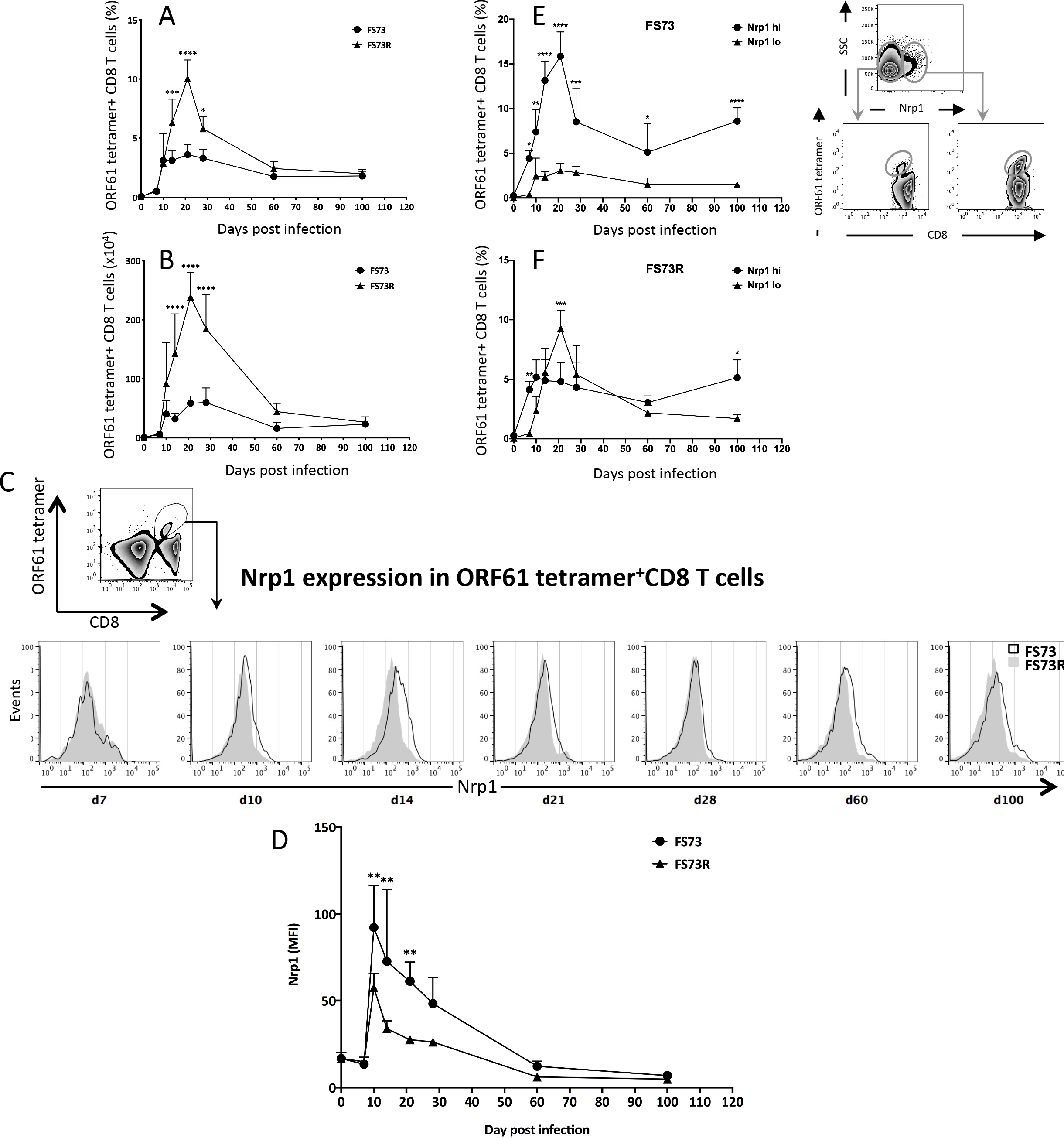
Nrp1 expression on CD8 T cells after persistent (FS73R) and non-persistent (FS73) MHV-68 infection. (A) The proportion of ORF61-specific T cells among total splenic CD8 T cells after infection with either the FS73 or FS73R strain of MHV-68. (B) Numbers of ORF61-specific CD8 T cells in spleens of mice infected with either the FS73 or FS73R strain of MHV-68. (C) Histograms showing Nrp1 expression gated on CD8^+^ ORF61 tetramer^+^ splenocytes at the times post infection shown. Y axes in bottom plots are normalized to the mode. (D) Nrp1 MFI of tetramer positive CD8 T cells compared over time for FS73 and FS73R infection. (E and F) Frequency of ORF61 tetramer^+^ CD8 T cells that were Nrp1^−^ or Nrp1^+^ after FS73 infection (E, panels on right show representative plots showing gating strategy to distinguish Nrp1^hi^ and Nrp1^lo^ cells) or FS73R infection (F). All data show mean ± SD of 4-5 mice per group; \*\**P*<0.01. Representative data from at least two experiments are shown.

### Tamoxifen-induced Nrp1 excision and YFP expression on CD8 T cells

In order to determine the role of Nrp1 in the antiviral CD8 T cell response, we wished to use an inducible knockout system that deletes Nrp1 selectively in CD8 T cells, and only after induction by tamoxifen, as Nrp1 is important in the development of embryonic blood vessels and a systemic knockout is lethal^4^. We therefore exploited conditional Nrp1 knockout transgenic mice (Nrp1 E8i-CreERT2 R26-YFP; Nrp1cKO mice, Fig. 2) where the E8i-creERT2 cassette confers CD8 specificity^18^, but cre is translocated to the nucleus only after tamoxifen treatment. These mice also contain floxed Nrp-1 allele and a Rosa26-flox-stop-flox-YFP sequences resulting in deletion of Nrp-1 and expression of YFP in cells where cre is active. Then we characterized these mice to verify inducible deletion by treating Nrp1cKO mice or B6 with tamoxifen (1 mg/mouse) for 5 consecutive days (Fig. 3A), and measured Nrp1 expression on CD8 T cells 48 hours after the last treatment. We observed that YFP expression was induced on CD8 T cells in Nrp1cKO (Fig. 3B, third panel) but not B6 mice (Fig. 3B, second panel). While there was only a very small population of CD8 T cells expressing YFP after vehicle treatment (Fig. 3B, first panel), this rose to 66% following tamoxifen treatment (Fig. 3B, third panel). Tamoxifen-mediated induction of YFP was not observed in CD4 T cells in these mice (Fig. 3B, forth panel), confirming that cre-mediated deletion was limited to CD8 T cells. To confirm YFP expression correlated with Nrp1 cell surface expression, we stained for Nrp1 on CD8 T cells from mice treated as described, and found this molecule was absent from the YFP^+^ population, but present on a proportion of YFP^−^ cells (Fig. 3B, lower panels, Fig. 3C). After tamoxifen treatment there was still an easily detectable population of Nrp1^+^ CD4 T cells (Figs. 3B and 3C), demonstrating the absence of Nrp1 deletion in this population. These data confirmed that tamoxifen treatment effectively abrogated Nrp1 expression on CD8 T cells and cells lacking Nrp1 were marked by YFP fluorescence.

**Figure 2.**
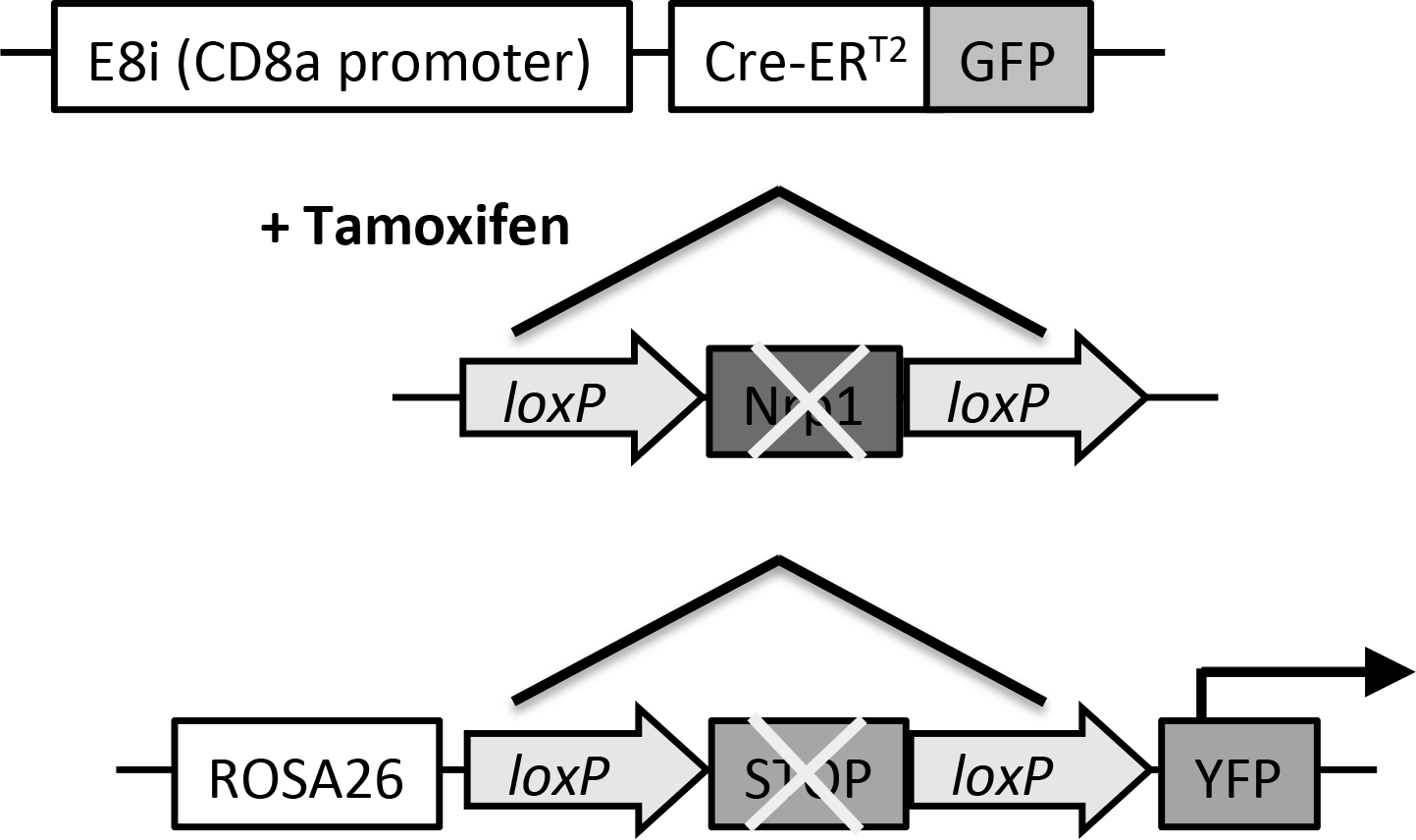
Schematic representation of Nrp1 E8i-CreERT2 R26-YFP mice. In the presence of tamoxifen, regions flanked by loxP sequences are excised, which deletes Nrp1 in CD8 T cells, and removes the stop sequence downstream of the ROSA26 promoter, resulting in YFP expression on CD8 T cells.

**Figure 3.**
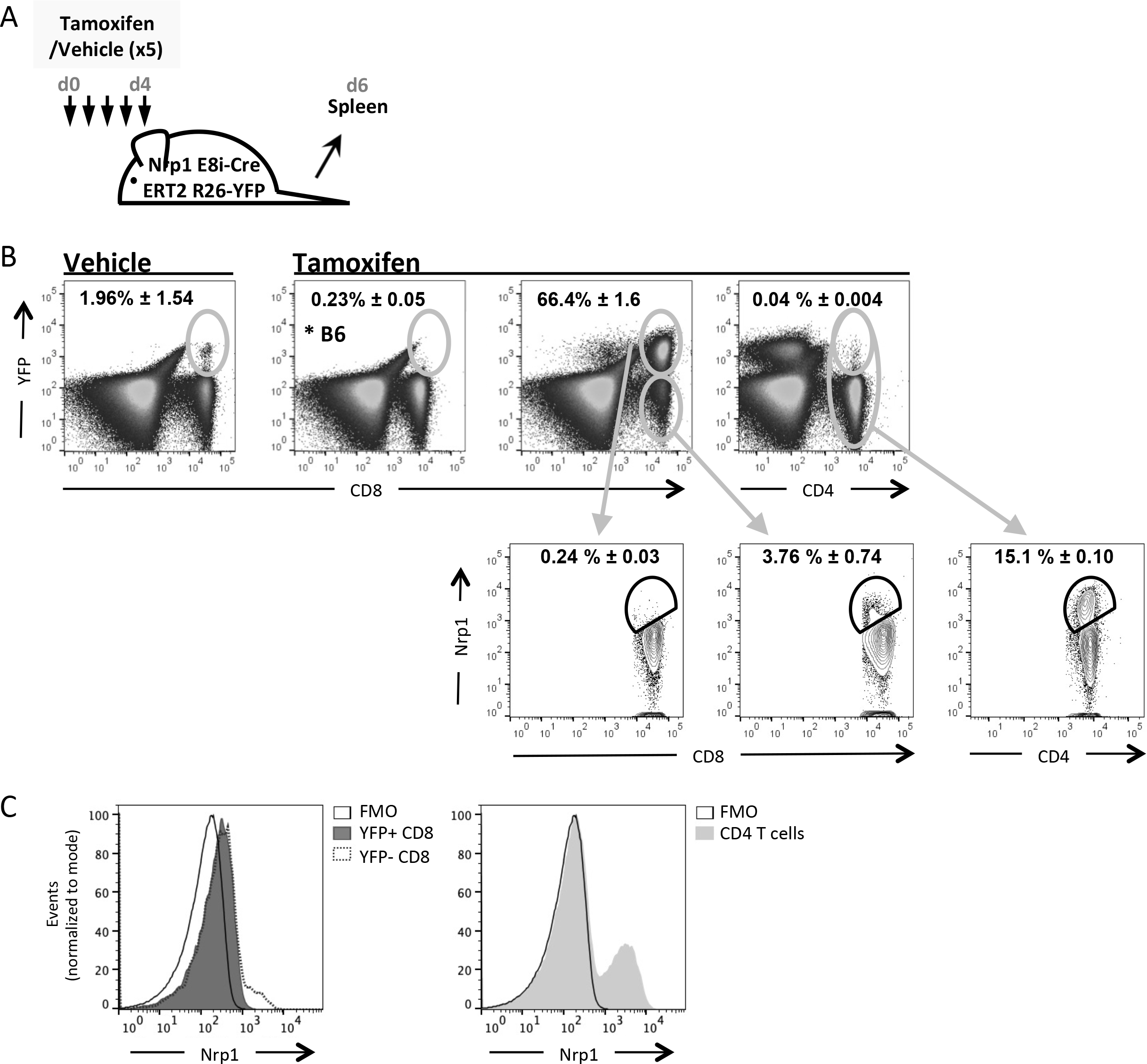
Tamoxifen administration effectively deltes Nrp1 from CD8 T cells in E8i-CreERT2 R26-YFP mice. (A) Protocol for tamoxifen treatment. (B) YFP expression was measured on CD8 or CD4 T cells after either tamoxifen or vehicle treatment in B6 or E8i-CreERT2 R26-YFP mice. Lower panels show Nrp1 staining on the indicated populations. (C) Histograms showing Nrp1 staining on CD8 (left) and CD4 (right) T cells from tamoxifen treated E8i-CreERT2 R26-YFP mice. Representative data from at least two experiments are shown. Percentages shown are among the total CD8 or CD4 population, as appropriate.

### Effect on the CD8 T cell responses of Nrp1 deletion

To test the effect of Nrp1 on CD8 T cell differentiation during murine gammaherpesvirus infection, we treated Nrp1 mice with tamoxifen as described above, and infected them with FS73 or FS73R two days after the last tamoxifen treatment (Fig. 4A). Vehicle treated mice exhibited a very small YFP+ population, but this was greatly enlarged after tamoxifen treatment (Fig. 4B). To measure the effect of Nrp1 on CD8 T cell expansion and memory formation, we compared the proportion of YFP^+^ and YFP^−^ cells that stained with a tetramer identifying CD8 T cells recognizing the dominant ORF61 epitope. In this way we had an internal control in each mouse, normalizing for variations in virus titers or other variables from mouse to mouse, and enabling the use of paired statistical tests for significance. The frequency of tetramer positive cells were not significantly different in YFP^+^ and YFP^−^ cells at 14 days post-infection, regardless of virus strain (Fig. 4C), indicating the magnitude of the effector response was not altered by absence of Nrp1. However we did detect differences in the differentiation status of the CD8 T cells. On day 14 in blood, we found that most tetramer^+^YFP^+^CD8 T cells (Nrp1 deleted) had the phenotype of precursors of memory cells (KLRG-1^−^CD127^+^), whereas the majority of tetramer^+^YFP^−^CD8 T cells (expressing Nrp1) were terminally differentiated effector T cells (KLRG-1^+^CD127^−^; Fig. 4D and 4E). These results were comparable between persistent and non-persistent infections, although in FS73R infection the KLRG-1^+^CD127^+^ population was as prominent as the KLRG-1^−^CD127^+^ population (Fig. 4E). We have previously shown that most memory CD8 T cells after infection with either the FS73 or FS73R viruses do not upregulate CD127, unlike that seen in other infection models^16^. Here we observed most YFP^+^ and YFP^−^ CD8 T cells in the spleen became KLRG-1^+^CD127^−^ by day 28 (Fig. 4F and 4G), indicating the absence of Nrp-1 does not affect this phenotype. These data showed Nrp1 favored the differentiation of effector CD8 T cells during the peak response, and in it’s absence T cells differentiated preferentially toward the memory precursor phenotype. However these changes did not endure to the memory phase, where both Nrp1 sufficient and deficient CD8 T cells had similar phenotypes.

**Figure 4.**
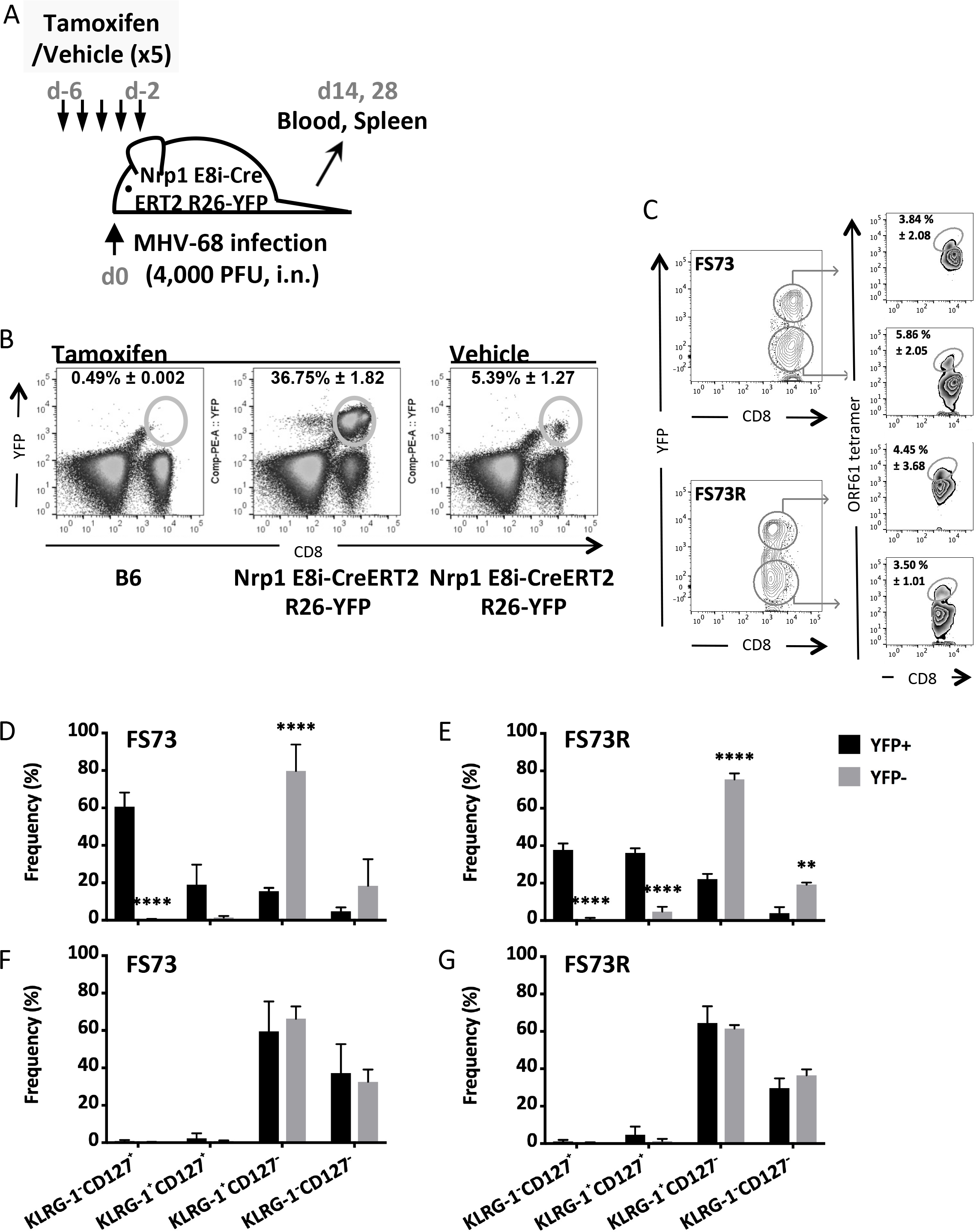
Effect of Nrp1 deletion on CD8 T cell responses. (A) Experimental design. (B) YFP expression in tamoxifen treated B6 and tamoxifen or vehicle treated E8i-CreERT2 R26-YFP mice. (C) Evaluation of the size of the ORF61-specific population in the blood within either the YFP^+^ or YFP^−^ CD8 T cell populations in FS73 or FS73R infected mice. (D-E) At d14 post-infection blood was stained to identify CD8^+^ORF61 tetramer^+^ cells, and the frequencies among this population are shown with respect to KLRG-1 and CD127 expression. (F-G) Spleen cells at d28 post-infection were stained as in (D-E). All data show mean ± SD of 4-5 mice per group; \*\**P*<0.01, \*\*\*\**P*<0.0001. Representative data from at least two experiments.

### Effect of Nrp1 on the recall response

Having observed a role for Nrp1 in the formation of memory CD8 T cells during the effector response, we next tested whether the quality of the resulting memory population was altered by re-challenging with virus. As in previous experiments, tamoxifen was administered before infection to delete Nrp1, then splenic CD8 T cells were purified on d28 post-infection, and adoptively transferred into congenic recipient mice (Fig. 5A). YFP expression was then used to identify the response from CD8 T cells with intact Nrp1 (YFP^−^) or deleted Nrp1 (YFP^+^; Fig. 5B). Recipients were rechallenged with virus, then six days later spleens were removed and the expansion of adoptively transferred cells measured by flow cytometry. Memory CD8 T cells from mice infected with FS73 expanded comparably regardless of Nrp1 expression status (Fig. 5C), whereas the expansion of memory cells from FS73R donors was significantly reduced when they lacked Nrp1 (YFP^+^, Fig. 5D).

**Figure 5.**
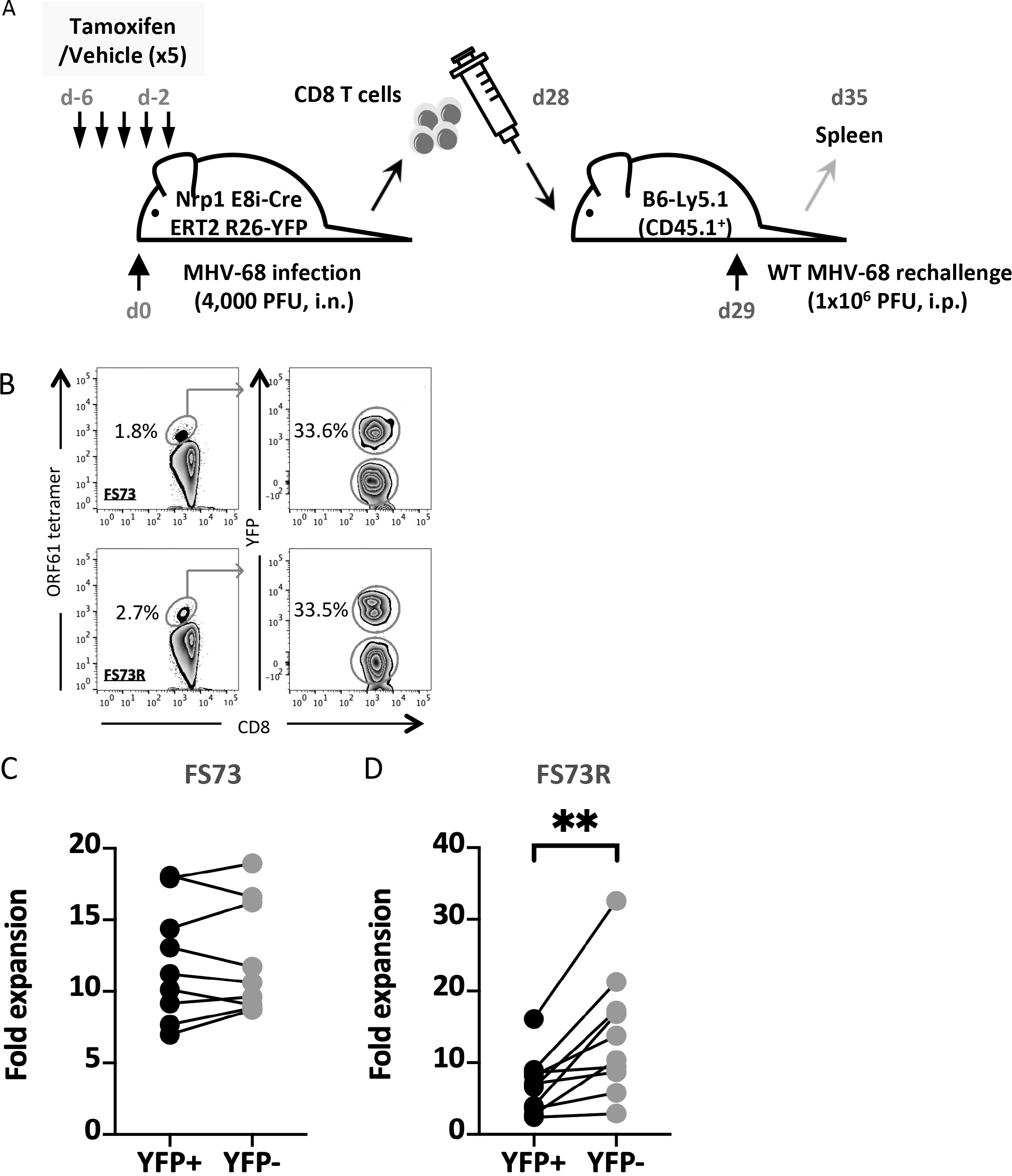
Effect on the recall response of Nrp1 deletion before infection. (A) Experimental design showing Nrp1 deletion before infection, then adoptive transfer of memory CD8 T cells followed by infection of congenic secondary hosts. (B) Flow cytometry plots of adoptively transferred CD8 T cell populations. Plots show ORF61 tetramer positive CD8 T cell populations (left) and YFP positive populations within the tetramer positive population (right) from FS73 (top) or FS73R (bottom) infected mice. (C-D) Graphs showing expansion of YFP^+^ and YFP^−^ CD8^+^ ORF61 tetramer^+^ populations from mice infected originally with either (C) FS73 or (D) FS73R after secondary exposure to WT MHV-68. Joined lines represented paired samples (YFP^+^ and YFP^−^ cells from the same mouse). Data combined from 2 experiments are presented in C and D. \*\**P*<0.01.

In the previous experiment Nrp1 was absent both during the primary CD8 T cell response, where memory ‘programming’ occurs, and also after CD8 T cells differentiated into memory cells. Next we tested whether the absence of Nrp1 during the secondary response alone affected T cell expansion. To examine this, we performed a similar experiment, but this time tamoxifen treatment started at d28 post-infection, then spleens were harvested two days after the cessation of treatment (Fig. 6A). As before, purified CD8 T cells were transferred into congenic recipient mice that were then infected with MHV-68. Six days later both YFP^+^ and YFP^−^ CD8 T cells from FS73 infected donors expanded to a similar extent (Fig. 6B). However YFP^+^ CD8 T cells from FS73R donor mice expanded to a greater extent than YFP^−^ cells (Fig. 6C), indicating Nrp1 expression during the secondary response limits the extent of T cell expansion. This greater expansion was attributed to better cell survival (lower frequencies of dead cells; Fig. 7A) and more actively proliferating cells (EdU incorporation; Fig. 7B) among the YFP^+^ population. These data indicated that Nrp1 restrains cellular proliferation and survival during the recall response, but it also plays a different role during the priming or post-priming phase of the response, promoting the development of memory cells capable of making an optimal recall response.

**Figure 6.**
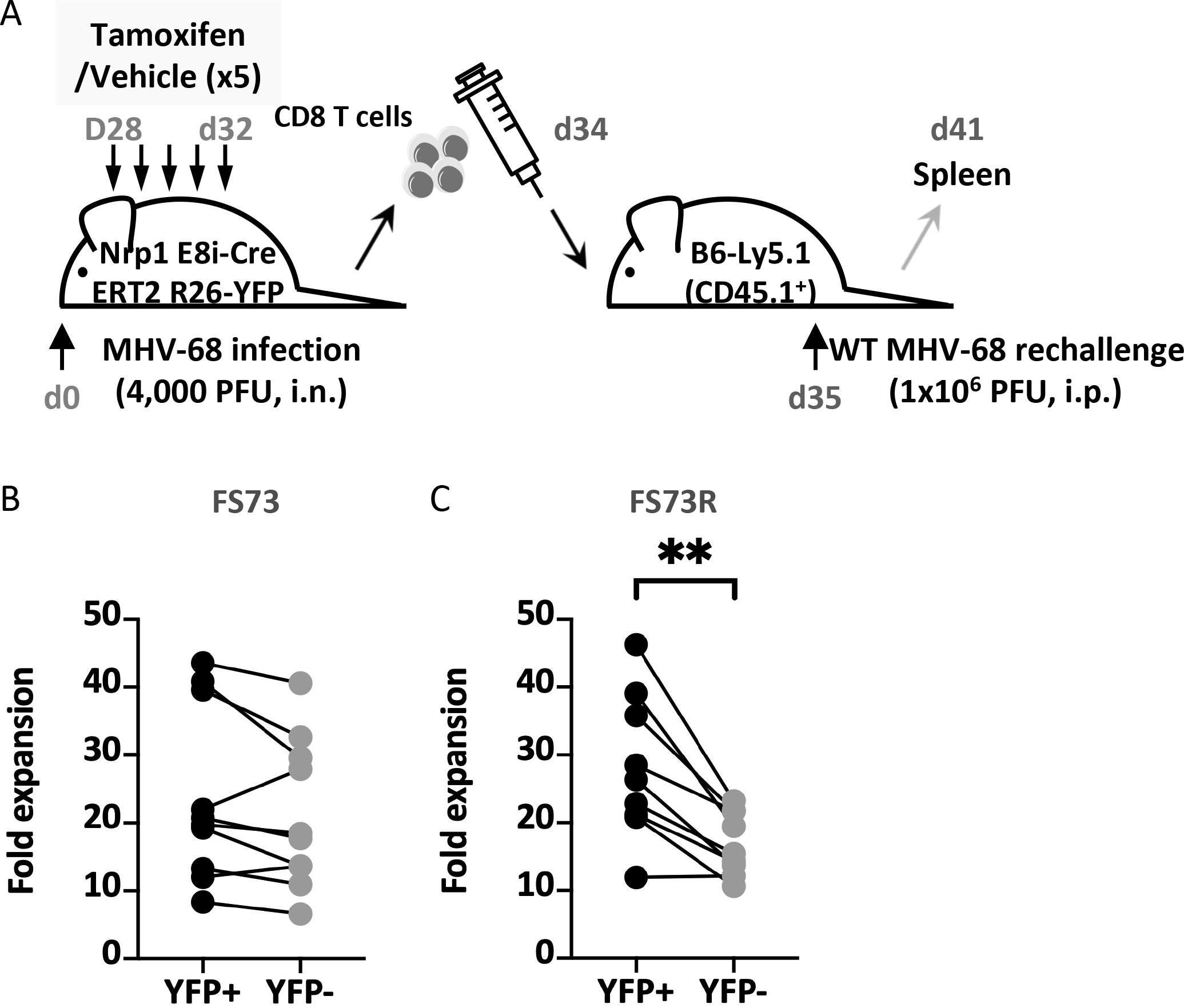
Effect of Nrp1 deletion just prior to the recall response. (A) Experimental design showing deletion of Nrp1 in memory cells, just before adoptive transfer to secondary congenic hosts. (B-C) Graphs showing expansion of YFP^+^ and YFP^−^ CD8^+^ ORF61 tetramer^+^ populations from mice infected originally with either (B) FS73 or (C) FS73R after secondary exposure to WT MHV-68. Joined lines represented paired samples (YFP^+^ and YFP^−^ cells from the same mouse). Data combined from 2 experiments are presented. \*\**P*<0.01.

**Figure 7.**
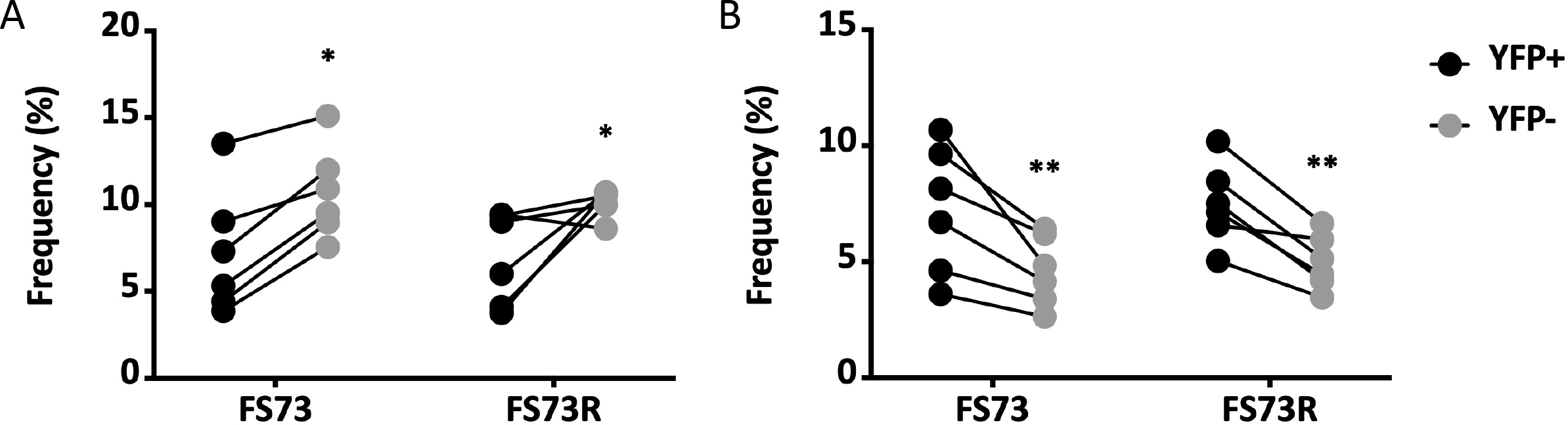
Cell viability and proliferation during recall responses where Nrp1 was deleted in memory cells. Experimental design was the same as that shown in Fig.6A. (A) Graph of the percentage of dead cells after the recall response, determined by gating on CD8^+^ ORF61 tetramer^+^ cells and a LIVE/DEAD™ stain. (B) Mice were injected with 100□l of 10□M Edu i.p. 16 hour before euthanasia. Edu incorporating proliferating cells were determined by gating on CD8^+^ ORF61 tetramer^+^ EdU^+^ cells after the recall response. All data show mean ± SD of 5-6 mice per group; \**P*<0.05, \*\**P*<0.01. Representative data from at least two experiments.

## DISCUSSION

This work shows clearly that neuropilin 1 plays an important role both in the differentiation of memory precursor cells and their capacity to mount a recall response. Cell surface Nrp1 expression was induced at the peak of the virus-specific CD8 T cell response, then declined slowly as these cells differentiated to memory cells. Interestingly we detected higher level induction in mice infected with the non-persistent FS73 strain. Our previous research showed a potent CD8 T cell response is induced after infection with either FS73 or FS73R strains of MHV-68^16^. However the capacity of the persistent strain to establish a latent infection in the spleen leads to splenomegaly, which increases the total number of virus-specific CD8 T cells, and also likely creates a more pro-inflammatory environment in the spleen. While the regulation of Nrp1 in T cells is not known, it is possible Nrp1 expression is restrained by these pro-inflammatory signals. It is particularly interesting that Nrp1 deletion only affects recall responses in FS73R infected mice, despite observing lower Nrp1 expression after infection with this persistent strain. The reasons for this effect are currently unclear, but may be due to the fact that during persistent infection there is sporadic reactivation, re-exposing the T cell response to viral antigens. This would lead to antigen presentation by infected B cells, which express the Nrp1 ligand semaphorin 4A^19^, potentially signaling to virus-specific memory CD8 T cells. In the acute infection with the non-persistent FS73 strain, these interactions with semaphorin 4A expressing antigen presenting cells would be limited to lung-draining dendritic cells and infected lung epithelial cells, but few B cells. Sporadic re-exposure to viral antigen during persistent infection endows the memory CD8 T cells with the ability to elaborate antiviral effector functions more quickly^16^, indicating a heightened state of readiness to counter the virus when it reactivates. Other evidence for these cells being in a different state of differentiation relative to memory cells in non-persistently infected animals includes lower levels of Bcl-2, lower IL-2 production and a faster turnover^16^. Maintaining memory CD8 T cells in this differentiation state may be more dependent on Nrp1, which may also have an impact on their ability to mount a recall response.

Our data show that, regardless of virus strain, the absence of Nrp1 on CD8 T cells at the time of infection resulted in a bias toward KLRG1^−^CD127^+^ memory precursor phenotype cells at the expense of KLRG1^+^CD127^−^ effector cells at the peak of the response. However this reduction in the proportion of effector cells did not reduce the overall magnitude of the response, indicating T cell proliferation is unaffected. Nrp1 therefore may have a role in restraining the differentiation of memory precursors, while favoring differentiation of effector T cells. This is counter to what may be expected based on a previous report that found Nrp1 at the immunological synapse acts through the phosphatase PTEN to restrain Akt phosphorylation in T_reg_^19^. In CD8 T cells Akt promotes effector cell differentiation and glycolytic metabolism acting through mTOR and Tbet^20^, whereas inhibition of Akt promotes memory differentiation^21,22^. Therefore Nrp1 may be expected to restrict Akt phosphorylation and promote memory differentiation, counter to that which we observed. This is likely due differences in the signal transduction pathways present in T_reg_ and CD8 T cells. Further work is necessary to uncover the underlying mechanism, and to determine why this apparent skewing to memory precursors does not result in a larger long-term memory population.

Our use of a conditional deletion model where Nrp1 expression is abrogated on approximately half the CD8 T cells, and the other half retain Nrp-1 expression, allowed us to perform very precise internally controlled experiments measuring the secondary CD8 T cell response. A complex picture of the roles of Nrp1 during the recall response emerges, with contrasting functions during different stages of the response. When deleted prior to infection, the absence of Nrp1 reduced the size of the recall response, indicating Nrp1 promotes the ability of memory cells to expand upon antigen re-exposure. While Nrp1 expression was most prominent on effector cells, it was still expressed at a low level on memory CD8 T cells at d28 post infection. Therefore Nrp1 could be acting in this context either by ‘programming’ appropriate differentiation of memory cells during the effector response, signaling during the contraction and early memory phases, or potentially both.

To interrogate the function of Nrp-1 during the recall response itself, we allowed the effector and memory response to develop in the presence of Nrp1 then deleted it just prior to memory cell harvest and cell transfer to secondary recipients. In this context deletion of Nrp1 enhanced the recall response, by promoting T cell viability and proliferation. This indicates during the differentiation of memory cells to effector cells Nrp1 serves to repress CD8 T cell proliferation and limit cell survival, presumably to limit the size of the T cell response and prevent immunopathology during the recall response.

A previous report detailed a role for Nrp1 in the initiation a primary T cell response from human T cells^23^. Antibody blockade of Nrp1 reduced the proliferation of naïve T cells 50-60% when stimulated with allogeneic dendritic cells, and Nrp1 was shown to cluster to dendritic cell:T cell contact areas. This contrasts with our finding that the primary response was not affected by the presence or absence of Nrp1. There are many differences between this report and our study, including the use of human cells vs mice, studying whole T cell populations vs CD8 T cells, and studying allogeneic responses vs antigen-specific responses. One additional potential reason for the difference in our findings is that antibody blockade of Nrp1 at the cellular interface may result in steric hindrance of other important interactions, thereby reducing signaling through the immunologic synapse. This is not a concern in our studies, as we used genetic deletion to ablate Nrp1 expression.

Our finding that Nrp1 expressed during the secondary response represses the proliferative response can be seen as consistent with previous studies reporting inhibition of T cell responses by Nrp1. Semaphorin 3A (Sema3A), a physiological ligand of Nrp1^24,25,26^, inhibits *in vitro* DC-T cell interaction^27^ and tumor-T cell interaction^28^, and by blocking Sema3A, hence suppressing the downstream signaling involving Nrp1, T cell activation and proliferation were restored. However these studies used whole T cell populations, which include T_reg_, so the inhibition observed may have been due to T_reg_-mediated suppression, in which Nrp1 plays a key role^8,29,30^, rather than direct interactions with CD8 T cells.

A previous report identified Nrp1 upregulation on anergic mouse CD8 T cells, but Nrp-1 did not appear to play any role in the tolerant phenotype^31^. However to our knowledge our study is the first to interrogate Nrp1 function directly on an antigen-specific antiviral CD8 T cell response *in vivo*. It highlights complex roles for this molecule in memory CD8 T cell differentiation and the secondary immune response, which depend upon the stage of the response and the nature of the infection. While Nrp1-semaphorin interactions are known to have an important impact on anti-tumor immunity, this study shows additional roles in antiviral immunity, and the memory CD8 T cell response.

## ACKNOWLEDGEMENTS

This work was funded in part by NIH grants AI122854 and AI117268. We thank Dr. Dario Vignali for providing Nrp1 conditional knockout mice, and for useful discussions.

